# Motion extrapolation in the flash-lag effect depends on perceived, rather than physical speed

**DOI:** 10.1101/2021.03.22.436374

**Authors:** Jane Yook, Lysha Lee, Simone Vossel, Ralph Weidner, Hinze Hogendoorn

## Abstract

In the flash-lag effect (FLE), a flash in spatiotemporal alignment with a moving object is misperceived as lagging behind the moving object. One proposed explanation for this illusion is based on predictive motion extrapolation of trajectories. In this interpretation, the diverging effects of velocity on the perceived position of the moving object suggest that FLE might be based on the neural representation of perceived, rather than physical, velocity. By contrast, alternative explanations based on differential latency or temporal averaging would predict that the FLE does not rely on such a representation of perceived velocity. Here we examined whether the FLE is sensitive to illusory changes in perceived speed that result in changes to perceived velocity, while physical speed is constant. The perceived speed of the moving object was manipulated using revolving wedge stimuli with variable pattern textures (Experiment 1) and luminance contrast (Experiment 2). The motion extrapolation interpretation would predict that the changes in FLE magnitude should correspond to the changes in the perceived speed of the moving object. In the current study, two experiments demonstrated that perceived speed and FLE magnitude increased in the dynamic pattern relative to the static pattern conditions, and that the same effect was found in the low contrast compared to the high contrast conditions. These results showed that manipulations of texture and contrast that are known to alter judgments of perceived speed also modulate perceived position. We interpret this as a consequence of motion extrapolation mechanisms and discuss possible explanations for why we observed no cross-effect correlation.

## 1. Introduction

In the process of visual perception, delays are incurred as information encoded by the retina is transmitted to the visual cortex (Schmolesky et al., 1998). Neural processing subsequently takes time, so for a time-varying stimulus such as a moving object, its position information would be outdated when it is available in visual cortical areas (De Valois & De Valois, 1991). Because the object would continue moving in the physical world, a lag between its perceived and veridical position would be inevitable at any given time (Maunsell & Gibson, 1992). Intuitively, the impact of such delays should be significant when localizing moving objects, but despite this, we seem to be able to accurately pursue and interact with even fast-moving objects (Smeets & Brenner, 1995b). One proposed explanation is that neural delays are partly compensated perceptually by a process of motion extrapolation (Nijhawan, 1994, 2002; motor compensation is also observed in Kerzel & Gegenfurtner, 2003 and outlined in Nijhawan & Wu, 2009). The motion extrapolation hypothesis proposes that the brain continuously extrapolates the trajectory of a moving object to predict where it is now, as closely as possible to its veridical position (Cavanagh, 1997; Nijhawan & Wu, 2009).

A strong case for visual motion extrapolation has been made on the basis of motion-induced position shifts, many of which indicate a strong coupling between motion and position signals. For example, the Fröhlich effect describes an illusory forward shift in the starting point of the motion trajectory (Fröhlich, 1924), and the representational momentum phenomenon involves a forward displacement in the stopping point of a moving object (Freyd & Finke, 1984; Hubbard, 1995, 2005, 2018). Similarly, when an object moves into a blindspot, the final position of the occluded object is extrapolated in the direction of motion beyond its vanishing point (Maus & Nijhawan, 2008); or if a moving object contains a moving texture, the perceived position of the object appears shifted toward the direction of motion of the texture, as opposed to motion of the object itself (Arnold, Thompson, & Johnston, 2007; De Valois & De Valois, 1991; Roach, McGraw, & Johnston, 2011). Of these illusions, this paper is concerned with the related flash-lag effect (FLE). The FLE has received extensive attention since Nijhawan (1994) reported that a flashed object is misperceived as lagging behind a physically aligned, continuously moving object. Explanations based on motion extrapolation mechanisms propose that the perceived position of the moving object is extrapolated, evoking an apparent spatial offset even though the two objects are always physically aligned (Nijhawan, 1994, 2002). Although different possible causes have been proposed for the FLE (see Hubbard, 2014; Maus, Khurana, & Nijhawan, 2010; Nijhawan, 2002 for reviews), convergent evidence from neural, behavioral, and computational studies demonstrate how extrapolation could be implemented in the visual system – and that this could compensate for neural delays and potentially explain the above illusions (Hogendoorn, 2020). Were these mechanisms to underlie the FLE, then such a shift should be expected, indeed, for the moving object but not the flash (Maus & Nijhawan, 2006; Nijhawan, 2002).

According to the motion extrapolation interpretation, the predictive mechanism that results in the FLE and related illusions derives from a process of constantly estimating and updating motion and/or position signals (Kwon, Tadin, & Knill, 2015). Arguably, early extrapolation signals have been shown to carry an estimate of an object’s velocity, in order to gauge the distance for the object to (have) travel(ed) (Crick & Koch, 1995; Nijhawan, 2008; Nijhawan & Wu, 2009; Pollen, 2003; Rosenbaum, 1975). Importantly, in both representational momentum (Freyd & Finke, 1985; Hubbard & Bharucha, 1988) and the FLE (Brenner & Smeets, 2000; Krekelberg & Lappe, 1999, 2000; López-Moliner & Linares, 2006; Murakami, 2001; Nijhawan, 1994; Nijhawan, Watanabe, Khurana, & Shimojo, 2004), the perceived offset increases linearly with the velocity of the moving object. However, not all FLE studies are consistent. Kanai, Sheth, and Shimojo (2004) observed that the FLE was unaffected by velocity when FLE magnitudes was measured in spatial units and, in fact, decreased with increased velocity in units of time. Cantor and Schor (2007) showed that the FLE decreases for velocities greater than one degree of visual angle (hereafter denoted as dva throughout the manuscript) per second, while Wojtach, Sung, Truong, and Purves (2008) tested a larger range of 3 dva/s–50 dva/s and found that the FLE varied as a logarithmic function of velocity, so the linear effects consistent with previous observations were only shown at the lower velocity ranges. Such inconsistent effects led us to question whether these FLE findings might be due to changes in perceived, rather than physical velocity (Finke, Freyd, & Shyi, 1986; Makin, Stewart, & Poliakoff, 2009). Although physical velocity and perceived velocity are naturally closely correlated, the distinction is that (physical) velocity could be perceived differently depending on properties such as object contrast (Stone & Thompson, 1992), transiency/duration (Treue, Snowden, & Andersen, 1993), as well as spatial (Smith & Edgar, 1990) and temporal (Shen, Shimodaira & Ohashi, 2003) frequency. For this reason, we investigate the nature of the velocity representation that contributes to the neural computation that generates the perceived offset in the FLE.

In the present study, we test the hypothesis that the magnitude of the FLE depends on perceived velocity, resulting from changes in perceived speed, even when the physical velocity of the moving object is unchanged. Unique among explanations of the FLE, under the motion extrapolation model, a neural representation of velocity is necessary for an explicit computation that allows us to correctly perceive objects in their current and future positions despite neural delays. Other models of the FLE which attribute the FLE to temporal processes (based on differential processing latency and attentional shifts; Baldo & Klein, 1995; Purushothaman, Patel, Bedell, & Ögmen, 1998; Ögmen, Patel, Bedell, & Camuz, 2004; Whitney & Murakami, 1998, or based on temporal averaging of positions, sampling, and postdiction; Brenner & Smeets, 2000; Eagleman & Sejnowski, 2000, 2007; Krekelberg & Lappe, 1999, 2000; Whitney, Murakami, & Cavanagh, 2000) would also expect the FLE to increase with faster physical speed – but only as a temporal offset evident as a spatial error (Kanai et al., 2004; Krekelberg and Lappe, 1999, 2000). These models therefore do not require an explicit representation of velocity, and as such, would not be expected to depend on perceived speed when physical speed remains constant.

## 2. Experiment 1

One method of manipulating perceived speed is by dynamically modulating the texture of a moving object (Treue et al., 1993). When Carlson, Schrater, and He (2006) presented a smaller square with a static noise texture superimposed on top of a larger square with dynamic noise and the two objects moved together, the area of static noise was perceived to consistently lag behind the area of dynamic noise – causing an illusory percept that they called the Floating Square illusion. Despite having the same physical velocities, observers perceived the dynamic noise object to move faster and separate from the static noise object. The difference in perceived speed of the two patterns was sufficient to generate large perceptual errors when their relative positions were in fact identical. The results from this experiment are consistent with the idea that motion extrapolation is influenced by perceived speed. We apply this manipulation to an FLE paradigm, where our motion stimulus contains either a static or a dynamic pattern.

Participants carried out a two-alternative forced-choice (2AFC) spatial localization task where they were asked whether a revolving stimulus or a flashed target appeared to lead in the direction of motion, at the moment of the flash. Hereafter we refer to this as the flash-lag task (Experiment 1A). In a control speed-discrimination task (Experiment 1B), participants were asked whether the patterns differed in perceived speed. We hypothesized that if the FLE were sensitive to a change in perceived speed, then both perceived speed and perceived offset would be greater for the dynamic stimulus relative to the static stimulus. Conversely, if the FLE were not dependent on perceived speed, then we would expect no difference in FLE magnitudes as an addition of a dynamic pattern.

### 2.1 Participants

A total of 99 observers participated in one or both parts of Experiment 1: 30 observers participated only in Experiment 1A, and 20 observers participated only in Experiment 1B. To investigate possible correlations between the FLE measure obtained in Experiment 1A and the perceived speed measure obtained in Experiment 1B, a further 49 observers participated in both tasks. Two observers were excluded from Experiment 1A because their results significantly violated normality (Shapiro-Wilk: *p* < 0.001; Kolmogorov-Smirnov: *p* < 0.001). One observer was excluded from both tasks for incorrectly performing the tasks. Together, this yielded a total of 76 observers in Experiment 1A (23 males, mean age 24.2 years and SD 6 years) and 68 observers in Experiment 1B (23 males, mean age 24.2 years and SD 5.6 years), with 48 of these observers having completed both tasks.

All observers were naïve to the purpose of the experiment, were right-handed, and reported normal or corrected-to-normal vision. Observers gave informed consent and were reimbursed 10 AUD per hour for their participation in the experiment. The study was conducted in accordance with the Declaration of Helsinki and approved by the University of Melbourne Human Research Ethics Committee of the Melbourne School of Psychological Sciences (Ethics ID 1954146.2).

### 2.2 Stimuli

All stimuli were presented on a 24.5-inch ASUS ROG PG258 monitor (ASUS, Taipei, Taiwan) with a resolution of 1920 × 1080 running at 200 Hz, controlled by an HP EliteDesk 800 G3 TWR running MATLAB R2017b (Mathworks, Natick, NA) with PsychToolbox 3.04.14 extensions (Kleiner et al., 2007). Participants sat in a light-attenuated room and viewed the stimuli from a headrest at 60 cm from the screen.

The motion stimulus consisted of a wedge segment superimposed on a static annulus, which was centered at a white fixation point and displayed on a uniform 50 % gray background (Figure 1A). The wedge’s inner and outer edges were 4.05 dva and 6.75 dva away from the fixation point and subtended 45 degrees of polar angle (hereafter denoted as ° throughout the manuscript; whereas, dva denotes degrees of visual angle) along the radial axis. The wedge revolved at a fixed speed of 200 °/s. The direction of motion varied randomly between clockwise or counter-clockwise across trials but was constant within a given trial.

**Figure 1.**
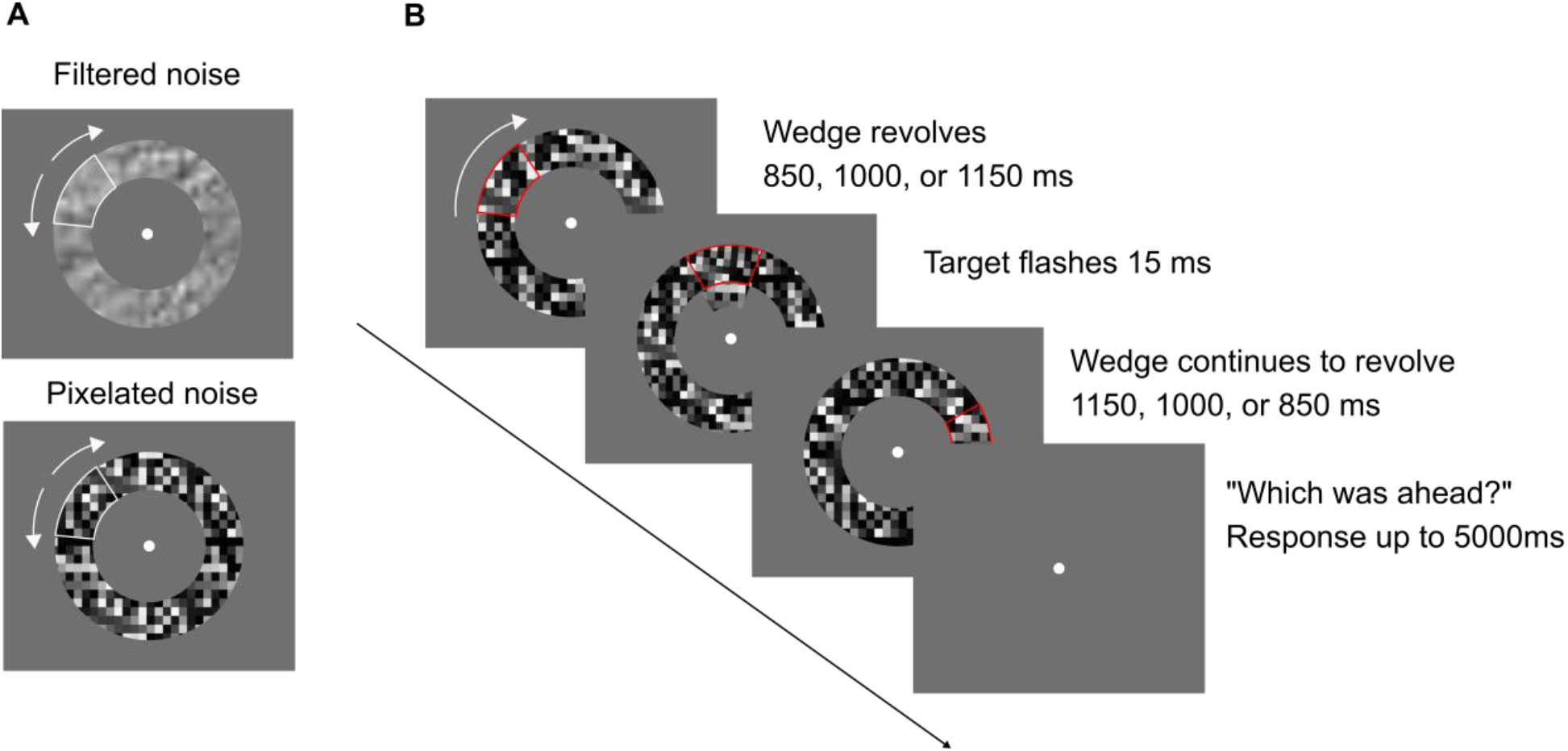
Schematic illustration of the stimulus configuration and flash-lag sequence in Experiment 1A. (A) Wedge and annulus containing filtered noise (top) and pixelated noise (bottom) texture (see also <u>videos</u> of the stimuli used in this experiment; speed reduced for illustrative purposes). (B) During a given trial, the wedge revolved within the annulus (shown with a pixelated noise texture and outlined in red in the figure). The target flashed after half the time of the overall sequence, and after a delay, observers were asked to respond using a keypress. A complete trial lasted up to 7 s, with the presentation sequence lasting 2 s. In Experiment 2A, the motion stimulus was a wedge segment without the annulus.

The wedge and annulus contained one of two textures. Filtered noise was created by applying a low-pass filter on white noise at 0.5 cycles/deg, and pixelated noise was created using an array of 5 × 5 pixel blocks with random luminance. The dynamic pattern was added by replacing the textures with a new random set of pixels, of no correlation with the previous set, at every frame. This configuration resulted in a flickering appearance of the texture. The background of the annulus always had the static pattern, because otherwise the wedge would be indistinguishable from the annulus if both textures were patterned dynamically (second-order motion).

In Experiment 1A, observers compared the position of the wedge to a target. The target was a stationary wedge with the same static texture (Figure 1B). Its inner and outer edges corresponded to 2.43 dva and 3.78 dva from fixation and subtended 45 ° along the radial axis.

In Experiment 1B, observers compared the speed of the wedge to a comparison. The comparison was a solid black wedge. The speed of the wedge was constant at 200 °/s, but the speed of the comparison varied across trials using a staircase procedure (see *2.3 Procedure* below). Both stimuli always revolved in the same direction, which was randomly chosen at the start of each trial.

### 2.3 Procedure

#### 2.3.1 Experiment 1A: Flash-lag task

On each trial, the wedge was presented from a random starting position of the annulus and revolved for one of three possible durations (850, 1000, or 1150 ms chosen at random). After half the time of the overall sequence, the target was presented for 15 ms along the inner circumference of the annulus, after which the wedge continued revolving for the remaining 1150, 1000, or 850 ms of its trajectory (cumulative presentation duration of 2 s). Then, observers were instructed to report whether the wedge or target was spatially ahead of the other when the target had appeared (2AFC). Keypress responses were recorded in the next 5 s – ‘i’ as in inner (target) or ‘o’ as in outer (wedge) – and prompted the next trial.

The relative spatial location of the wedge and target was driven by a one-up, one-down staircase procedure, and trials were randomly drawn from two interleaved staircases. At the first trial of each condition block, one staircase started at a large difference of +20 ° (moving ahead of flash). A –20 ° starting point was omitted in this procedure as a reversing, flash-lead presentation would be redundant for the purpose of our hypothesized flash-lag illusion. Instead, the second staircase started at a smaller difference of 0 ° (physically aligned). Depending on the response, the difference was adjusted by ±2 ° in the following trial drawn from that staircase. Observers completed 120 trials from each staircase, and the final 60 points of each staircase were averaged to estimate the point of subjective equality (PSE): the displacement at which the wedge and target were perceived as aligned.

Trials were blocked into 4 conditions. Each condition consisted of 120 trials (each lasting up to 7 s), for a total of 480 trials over 4 blocks. The conditions were determined based on a combination of *pattern type* (static or dynamic) and *texture type* (filtered noise or pixelated noise). The order of condition was randomized for observers who completed only Experiment 1A and counterbalanced if they completed both Experiment 1A and 1B. All observers completed 12 trials as practice before each experimental block, followed by a self-paced break. This resulted in a total testing duration of approximately 30 minutes.

#### 2.3.2 Experiment 1B: Speed-discrimination task

On each trial, the comparison was presented from a random starting position against a 50 % gray background and revolved for a randomly chosen duration of 1500, 1750, or 2000 ms (Figure 2).

**Figure 2.**
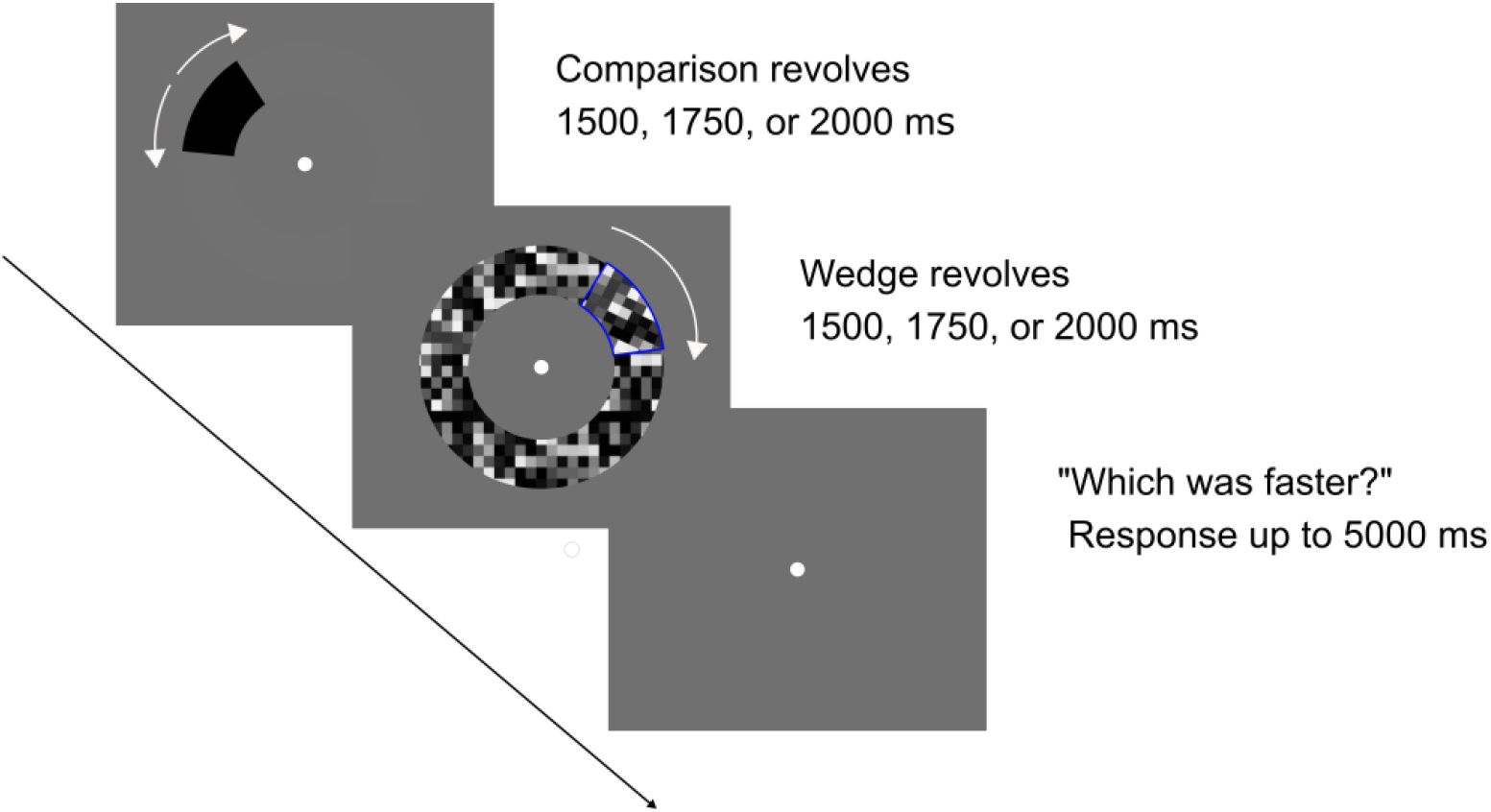
Schematic illustration of Experiment 1B. During a given trial, the comparison and the wedge within the annulus (shown with a pixelated noise texture and outlined in blue in the figure) were presented sequentially. After a delay, observers were asked to respond using a keypress. A complete trial lasted up to 9 s, with the presentation sequence lasting up to 4 s. In Experiment 2B, the motion stimulus was a wedge segment without the annulus.

Then, the comparison was removed from display, and the wedge from Experiment 1A was presented from a random starting position of the annulus and revolved for 1500, 1750, or 2000 ms also randomly chosen. Presentation duration and starting position were varied so that observers could not reliably base their responses on the difference between the distance traveled by each wedge. After the sequence ended, observers were instructed to report whether the comparison or the wedge revolved faster (2AFC). Keypress responses were recorded in the next 5 s – ‘1’ for first (comparison) or ‘2’ for second (wedge) – and prompted the next trial.

While the speed of the wedge was fixed, the speed of the comparison was determined using a similar staircase procedure using the ratio of these two stimulus speeds. Trials were drawn from one of two interleaved staircases. The first staircase was initialized at 1.075^7^ °, and the second started at 1.075^−7^ °, with a step size set to 1.075 ° for both. The staircase value of a given trial was proportionate to the relative speed of the comparison and the wedge. Because velocity is represented logarithmically in perception (Nover, Anderson, & DeAngelis, 2005; Priebe & Lisberger, 2004), the steps were adjusted by means of multiplying and dividing, rather than adding and subtracting. For example, 1.075^7^ ° (~ 1.66) indicated that the comparison revolved at 166 % of the speed of the wedge (166 % of 200 °/s). Depending on the response, the speed of the comparison was adjusted by 1.075 ° in the following trial drawn from that staircase. Observers completed 72 trials from each staircase, and the last 36 points of each staircase were averaged to estimate the PSE: the speed at which the comparison and wedge were perceived to match.

Trials were blocked into the same 4 conditions of Experiment 1A: *pattern type* (static or dynamic) and *texture type* (filtered noise or pixelated noise). All observers completed a total of 4 blocks. Observers who completed only Experiment 1B completed 72 trials per block (for a total of 288 trials), and observers who completed both Experiment 1A and 1B completed 144 trials per block (for a total of 576 trials). The total testing duration was 25 and 50 minutes, respectively.

### 2.4 Results

We tested for differences in the magnitude of the FLE and perceived speed in each condition using a two-way repeated measures ANOVA with factors of *pattern type* (static or dynamic) and *texture type* (filtered noise or pixelated noise). Post hoc paired-samples *t*-tests (two-tailed) with Bonferroni corrections estimated these differences.

#### 2.4.1 Experiment 1A: Effect of dynamic pattern on perceived flash-lag

A repeated measures ANOVA revealed a significant main effect of pattern type (*F*_1,75_ = 5.14, *p* = 0.026), with greater FLE magnitudes observed for dynamic pattern than for static pattern wedges (Figure 3A). There was no significant main effect of texture type (*F*_1,75_ = 1.61, *p* = 0.209) and no interaction effect between the two factors (*F*_1,75_ = 0.81, *p* = 0.37). The mean PSE was significantly larger for dynamic pattern compared to static pattern wedges for the filtered noise texture (*t*_75_ = 2.49, *p* = 0.015), but the difference did not reach significance for the pixelated noise texture (*t*_75_ = 1.03, *p* = 0.309).

**Figure 3.**
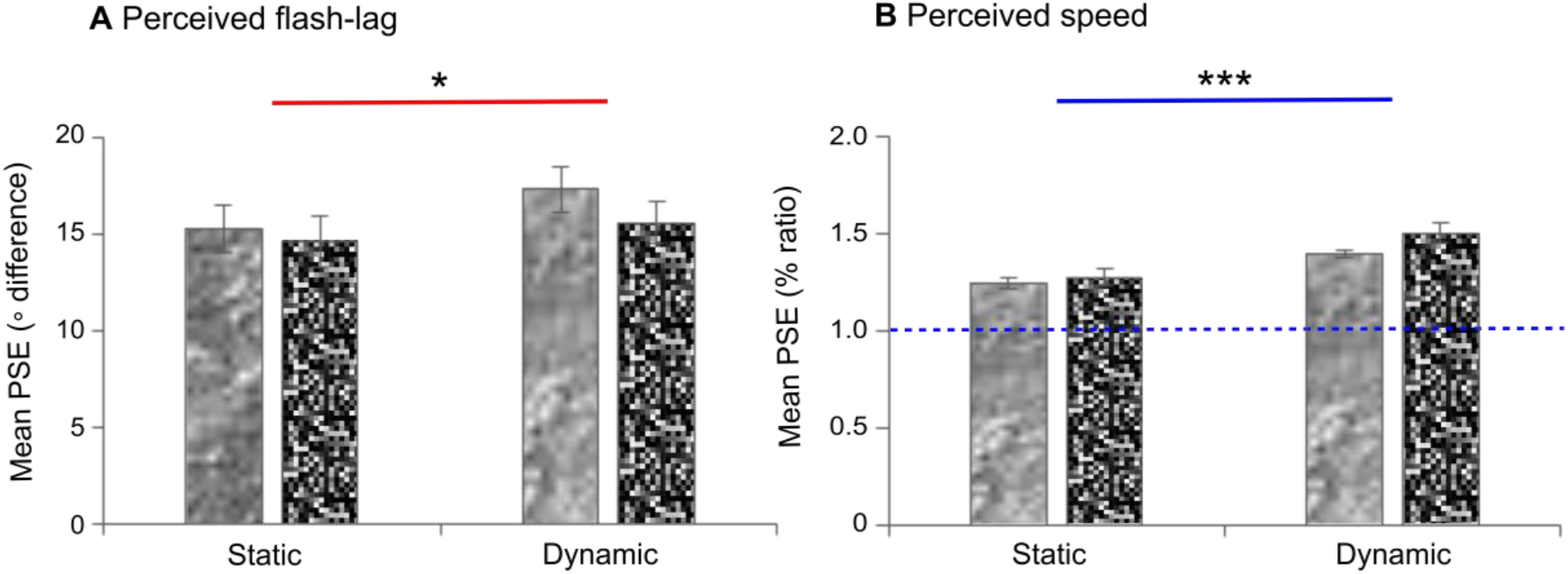
Results of Experiment 1. For illustrative purposes, the bars contain corresponding textures in the figure. (A) Mean FLE magnitudes of static pattern and dynamic pattern wedges across the texture conditions in Experiment 1A. (B) Mean perceived speeds of static pattern and dynamic pattern wedges across the texture conditions in Experiment 1B. Perceived speed was calculated as a ratio of the comparison speed relative to the wedge speed (taller bars mean that objects in that condition were perceived as faster). Baseline speed is indicated by a horizontal dashed line at y = 1. Mean perceived speeds were significantly different from zero and between pattern conditions. Error bars represent standard errors across observers. Asterisks above the bars indicate a statistically significant main effect of pattern type. * denotes *p* < 0.05. *** denotes *p* < 0.001.

#### 2.4.2 Experiment 1B: Effect of dynamic pattern on perceived speed

A repeated measures ANOVA showed a significant main effect of pattern type on perceived speed (*F*_1,67_ = 35.82, *p* < 0.001), with higher perceived speeds reported for dynamic pattern than for static pattern wedges (Figure 3B). This effect was significant for both filtered noise (*t*_67_ = 4.08, *p* < 0.001) and pixelated noise (*t*_67_ = 5.33, *p* < 0.001) textures. In addition, there was a significant main effect of texture type (*F*_1,67_= 11.5, *p* = 0.001) but no significant interaction effect (*F*_1,67_ = 2.518, *p* = 0.117).

#### 2.4.3 Cross-effect correlation

In order to further evaluate the possible relationship between the two measures, we investigated whether a dynamic pattern affected perceived speed and perceived position (flash-lag) in the same way. We were interested in the relation between one effect and the other, and if this was consistent between observers in a cross-effect correlation analysis. Among 48 observers who completed both tasks, we hypothesized that observers for whom the dynamic pattern textures greatly increased perceived speed would be expected to report larger changes in the illusion, and conversely, observers with minimal (or negative) changes in perceived speed would be expected to show corresponding small or negative effects in the illusion. However, Pearson’s correlation (two-tailed) did not reveal a significant correlation between the two tasks in either filtered noise (r = 0.15, *p* = 0.3) or pixelated noise (r = 0.07, *p* = 0.64) conditions (Figure 4).

**Figure 4.**
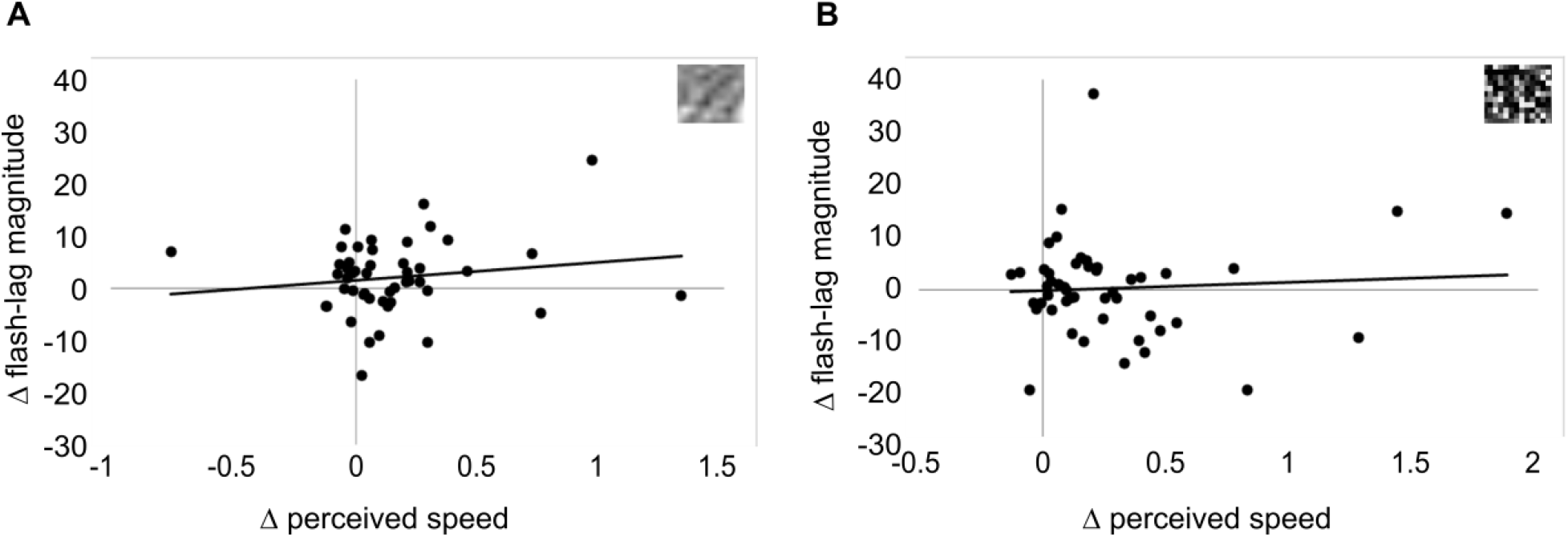
Change in FLE magnitude plotted as a function of change in perceived speed for (A) filtered noise and (B) pixelated noise textures. Legend for texture type is shown in the top right corner of each panel.

## 3. Experiment 2

Overall, Experiment 1 demonstrates that the dynamic modulation of a moving object’s texture, which is known to influence perceived speed, also influences the perceived position of the moving object – systemically in the same direction in the FLE, as predicted by the motion extrapolation model. To provide confidence that this effect is due to changes in perceived speed rather than unknown low-level factors, we carried out a parallel set of experiments to investigate whether a different, independent manipulation of perceived speed similarly affects the FLE.

The well-known ‘Thompson effect’ describes how speed is underestimated at low contrasts (Anstis, 2003; Blakemore & Snowden, 1999; Snowden, Stimpson, & Ruddle, 1998; Thompson, 1982). As a natural example, this has been used to explain why drivers frequently drive too fast under foggy conditions, compensating for their incorrect estimates of how fast (or slow) they and other cars are moving (Anstis, 2003; Snowden et al., 1998; but see also Owens, Wood, & Carberry, 2010; Pretto, Bresciani, Rainer, & Bülthoff, 2012). Theoretically, this effect is consistent with the findings of Berry, Brivanlou, Jordan, and Meister (1999), which showed that anticipatory neural responses for the leading edges of a moving object varied based on the contrast of the object. It also resembles the Hess effect (Hess, 1904), a similar illusion to the Floating Squares illusion and the FLE, in which a brighter object is perceived to lead a dimmer object when the two are actually aligned and have the same physical velocities. However, the effects of contrast on perceived speed depend on properties such as object luminance or physical velocity, and a handful of studies previously demonstrated that the effect can reverse under certain circumstances. For example, perceived speed can be overestimated at higher velocities (>8 dva/s; Hawken, Gegenfurtner, & Tang, 1994; Pretto et al., 2012; Thompson, Brooks, & Hammett, 2006) and low luminance levels (Vaziri-Pashkam & Cavanagh, 2008, 2011).

The effects of contrast have also been specifically investigated in the context of motion extrapolation. In the FLE, Kanai et al. (2004) found increases in FLE magnitude with decreases in contrast of moving objects with the background. Kanai et al. suggested that the FLE was modulated by positional uncertainty, as visibility is affected when the contrast of a target and background is reduced. Maus and Nijhawan (2006, 2009) also reported that the forward displacement of a moving object increases if the contrast of the target gradually decreased, while Hubbard and Ruppel (2014) reported that representational momentum decreases with low or decreasing contrast of the target and background. Finally, Vaziri-Pashkam and Cavanagh’s (2011) FLE paradigm presented a moving patch of random dots against the background containing static random dots. While they found that the perceived speed of moving objects increases at low luminance, caused by motion blurring for faster speeds (Vaziri-Pashkam & Cavanagh, 2008), there were no additional effects on FLE magnitude due to these changes in perceived speed.

We reexamined the effects of contrast in Experiment 2. In line with Experiment 1, the primary (moving) stimulus revolved at a speed of 200 °/s, but presented at either high (100 %) or low (10 %) contrast. Based on the divergent effects of contrast on perceived speed in the previous literature as briefly discussed above, we had no a priori directional hypothesis about the effect of contrast on perceived speed. Instead, we hypothesized that any effect of contrast on perceived speed would be mirrored in an effect on FLE magnitudes in the same direction. In other words, mean FLE magnitudes would increase in high contrast relative to low contrast conditions if perceived speed increased with high contrast, or in the opposite direction, it would increase for low contrast compared to high contrast conditions if perceived speed increased with low contrast. Conversely, if the FLE were not driven by perceived speed, then we would expect an effect of contrast on perceived speed, but no effect on FLE magnitudes.

### 3.1 Participants

Seven observers (3 males, mean age 25.3 years, SD 3.4 years) participated in the experiment. All were experienced observers and completed both Experiment 2A (Flash-lag) and 2B (Speed-discrimination). All inclusion criteria were identical to Experiment 1.

### 3.2 Stimuli

Stimuli were similar to those used in Experiment 1 with the following exceptions (see also <u>videos</u> of the stimuli used in this experiment). In Experiment 2, the motion stimulus consisted of a wedge segment without the annulus, against a uniform gray background. In doing so, we aimed to control for a potential confounding effect, as otherwise observers would be required to detect the contrast of the wedge on top of the background of the annulus against the background, while tracking the wedge’s motion (Blakemore & Snowden, 2000).

Observers viewed the stimuli from a distance of 50 cm. The inner and outer edges of the wedge and the comparison were 4.75 dva and 7.92 dva away from the fixation point and subtended 45 ° along the radial axis. The inner and outer edges of the target were 3.46 dva and 4.44 dva from fixation, subtending 45 ° along the radial axis. Stimuli textures were filtered noise or pixelated noise and always presented as a static pattern. The main experimental manipulation of Experiment 2 was that on each trial, the wedge was presented at high (100 %) or low (10 %) contrast relative to the gray background. The target was always presented at 100 % contrast.

### 3.3 Procedure

Due to the COVID-19 pandemic, observers participated from their own homes. All observers were provided with identical-speed ASUS ROG PG258Q monitors (1920 × 1080 resolution, 200 Hz), and completed the experiment in a dimly lit room. A chinrest was not used for this experiment, but observers were instructed to sit approximately 50 cm from the screen and make sure the fixation point was at eye level.

The experimental conditions were determined by a combination of *contrast level* (high or low) and *texture type* (filtered noise or pixelated noise). All observers completed a total of 4 blocks: 120 trials per block (for a total of 480 trials) in Experiment 2A and 144 trials per block (for a total of 576 trials) in Experiment 2B. The total testing duration was approximately 80 minutes.

### 3.4 Results

As in Experiment 1, we probed FLE magnitudes for each condition in Experiment 2A and perceived speed in Experiment 2B. We submitted all data to a two-way repeated measures ANOVA with factors of *contrast level* (high or low) and *texture type* (filtered noise or pixelated noise) and *t*-tests. All relevant assumptions were met.

#### 3.4.1 Experiment 2A: Effect of contrast on perceived flash-lag

A repeated measures ANOVA revealed a significant main effect of contrast level (*F*_1,6_ = 7.32, *p* = 0.035), with greater FLE magnitudes observed for low contrast than for high contrast wedges (Figure 5A). There was a main effect of texture type (*F*_1,6_ = 28.87, *p* = 0.002), with greater FLE magnitudes observed for pixelated noise than for filtered noise conditions. There was no significant interaction effect (*F*_1,6_ = 0.47, *p* = 0.52). The mean PSE was significantly larger for low contrast compared to high contrast wedges for filtered noise (*t*_6_ = 3.47, *p* = 0.013), but the difference did not reach significance for pixelated noise (*t*_6_ = 2.12, *p* = 0.078). For individual data, the FLE consistently increased in low contrast compared to high contrast conditions for all but two observers (P6 for filtered noise and P3 for pixelated noise; Figure 5B).

**Figure 5.**
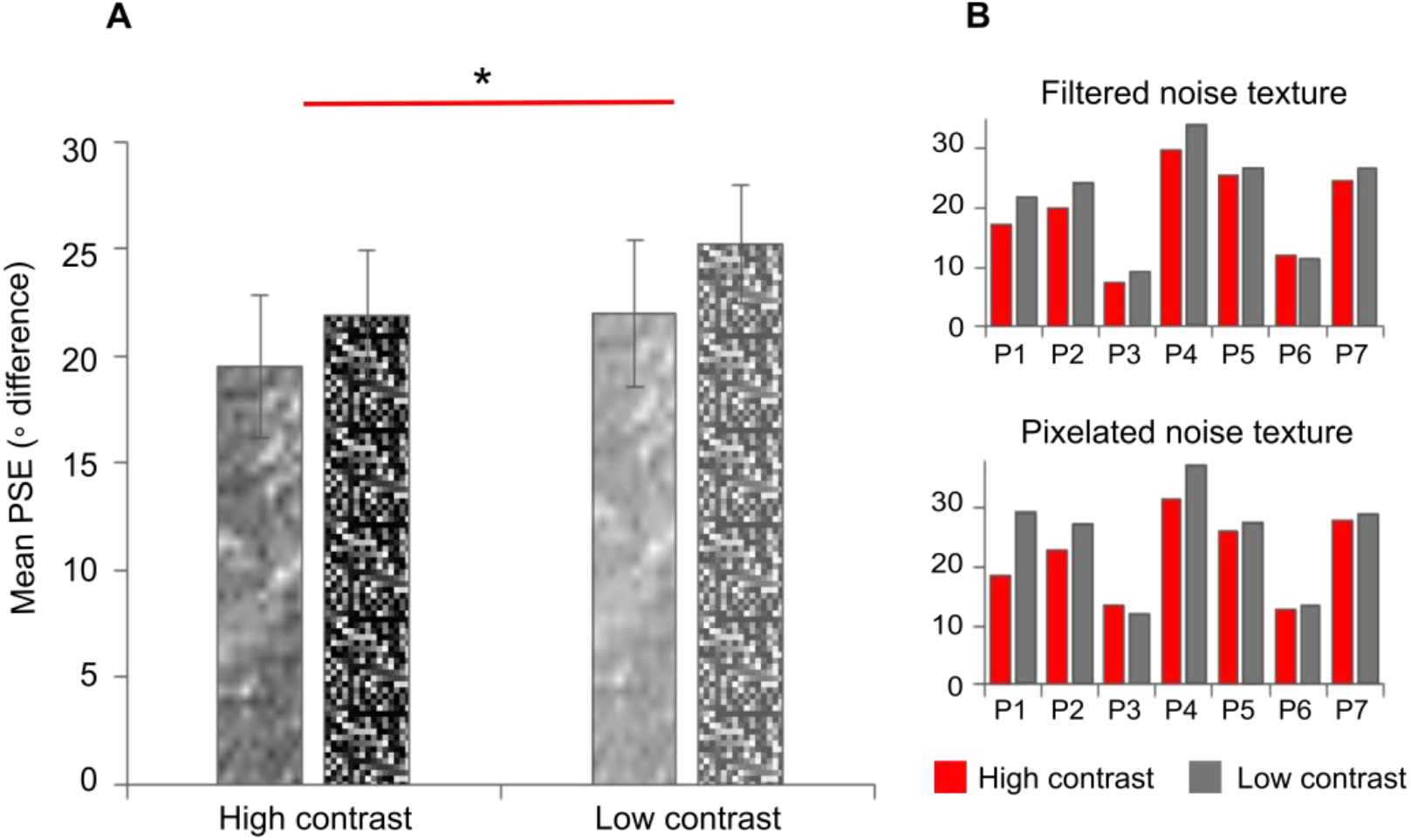
Results of Experiment 2A. For illustrative purposes, the bars contain corresponding textures and contrasts in this figure. (A) Mean FLE magnitudes of high and low contrast wedges across texture conditions, pooled over all seven observers. (B) Individual observers. Error bars represent standard errors across observers. The asterisk above the bars indicates a statistically significant main effect of contrast level at *p* < 0.05.

#### 3.4.2 Experiment 2B: Effect of contrast on perceived speed

A repeated measures ANOVA revealed a significant main effect of contrast level on perceived speed (*F*_1,6_ = 52.85, *p* < 0.001), with low contrast perceived to move faster than high contrast wedges of the same velocity (Figure 6A). There was no significant main effect of texture type (*F*_*1*,6_ = 0.086, *p* = 0.779) or an interaction effect (*F*_1,6_ = 4.68, *p* = 0.074). The mean PSE was significantly larger for low contrast compared to high contrast wedges in both filtered noise (*t*_6_ = 5.72, *p* = 0.001) and pixelated noise (*t*_6_ = 7.31, *p* < 0.001) conditions. The effect was highly consistent across individual observers (Figure 6B).

**Figure 6.**
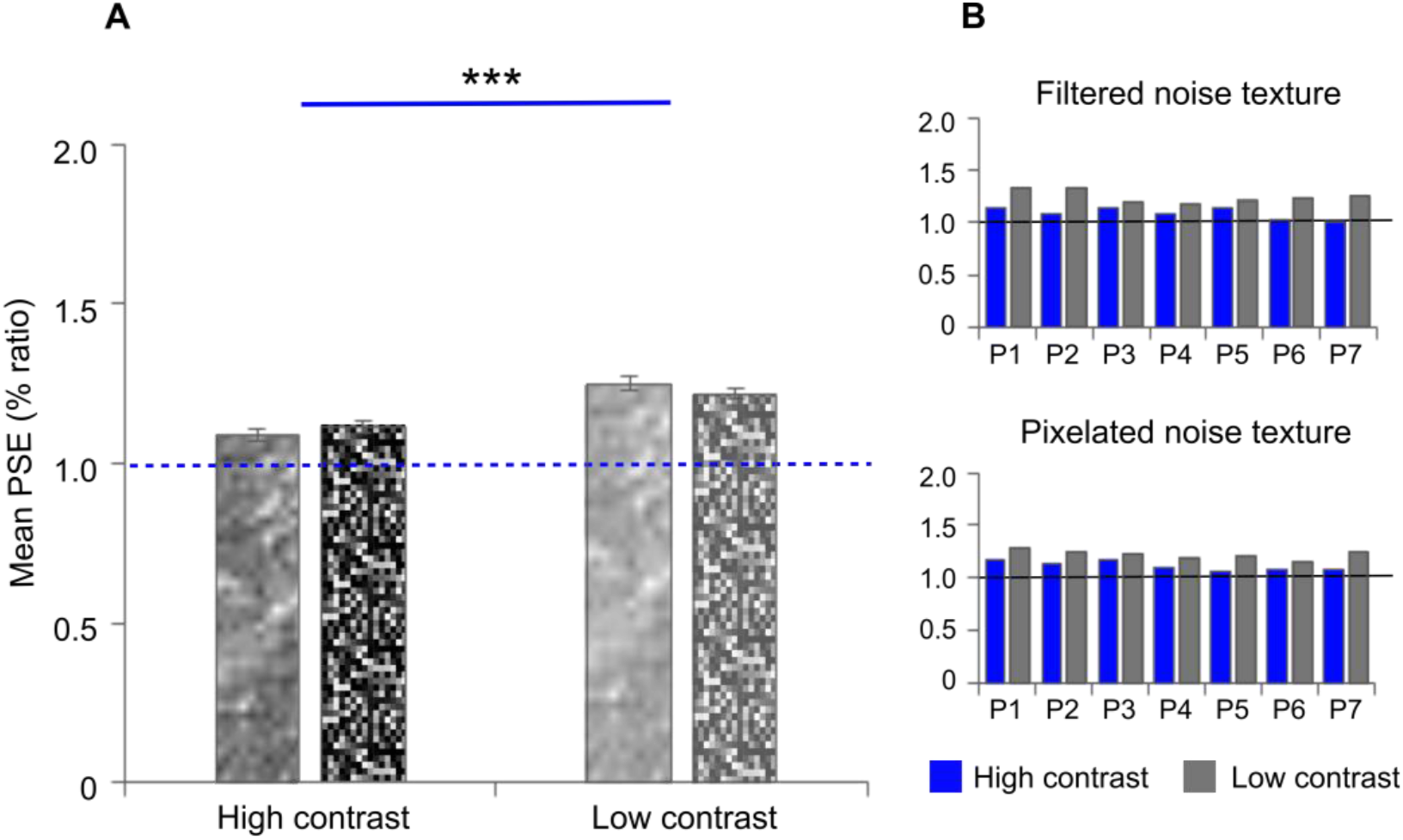
Results of Experiment 2B. For illustrative purposes, the bars contain corresponding textures and contrasts in this figure. (A) Mean perceived speeds of high and low contrast wedges across texture conditions, pooled over all seven observers. (B) Individual observers. Perceived speed was calculated as a ratio of the comparison relative to the wedge speed, and the baseline speed is indicated as y = 1. Error bars represent standard errors across observers. Asterisks above the bars indicate a statistically significant main effect of contrast level. *** denotes *p* < 0.001.

## 4. Discussion

The flash-lag effect (FLE) is a classic illusion that has been argued to result from motion extrapolation mechanisms that might be implemented in the visual system to contribute to overcoming its time delays. Although many alternative explanations for the FLE have been presented, the motion extrapolation account uniquely posits a neural computation that requires a representation of velocity. We hypothesized that under this interpretation, experimental manipulations that affect perceived velocity might similarly influence the magnitude of the FLE. Therefore, we examined how illusions of perceived speed could alter the perceived flash-lag in two experiments. We show that dynamically modulating a texture causes an illusory percept of increased speed as well as increased FLE magnitude compared to a static pattern. Similarly, when objects were presented at low contrast, both perceived speed and FLE magnitude also increased relative to high contrast. These findings appear to be consistent with the motion extrapolation account of the FLE.

Experiment 1 shows robust differences between dynamic and static patterns for perceived speed and FLE magnitude. Such differences were qualitatively similar across different textures. Overall, these results closely reproduce the previously reported effects on the perceived speed of gratings, Gabor patches, and in the Floating Square illusion (Carlson et al., 2006; De Valois & De Valois, 1991; Shen et al., 2003; Treue et al., 1993), and are also consistent with the effects of speed on motion extrapolation (Brenner & Smeets, 2000; Freyd & Finke, 1985; Hubbard & Bharucha, 1988; Krekelberg & Lappe, 1999, 2000; López-Moliner & Linares, 2006; Murakami, 2001; Nijhawan, 1994; Nijhawan et al., 2004).

Experiment 2 likewise shows subjective increases in both perceived speed and FLE magnitude at low contrast relative to high contrast. These results diverge from findings of previous motion extrapolation studies on the effects of contrast (e.g., Hubbard & Ruppel, 2014; Maus & Nijhawan, 2006, 2009), in that our motion stimulus was perceived as faster at low contrast than high contrast. The discrepancy with the Thompson effect is likely due to methodological differences because our wedge had revolved at approximately 28.15 dva/s, which is significantly faster than in the literature we outlined here (approximately 3-17 dva/s; Blakemore & Snowden, 1999; Hubbard & Ruppel, 2014; Maus & Nijhawan, 2006, 2009; Stone & Thompson, 1992; Thompson et al., 2006). Consistent with this interpretation, Vaziri-Pashkam and Cavanagh (2011) found that out of a range of velocities, only their highest velocity of 27.36 dva/s was overestimated at low contrasts.

If a neural representation of perceived velocity is used to inform the extrapolation process that underlies the FLE, then we would expect to see a causal relationship between the effects of dynamic pattern on perceived speed and the effects on the FLE. However, we did not observe a correlation between the behavioral responses, similar to Vaziri-Pashkam and Cavanagh (2011), with a portion of observers showing a decrease in FLE magnitude despite clear increases in perceived speed. One possibility is that the experiments in this study illustrate that different processes are involved in making judgments of speed versus relative position. For example, de’Sperati and Thornton (2019) showed that varying the contrast of a moving object influenced extrapolation judgments but not interceptive decisions when observers were required to make saccadic eye movements to avoid colliding with multiple moving targets. Based on these findings, de’ Sperati and Thornton suggested these tasks involve intrinsically different processes. An important question arising from our findings is therefore whether the representations of velocity that contribute to explicit reports of motion (i.e. *how fast* an object is perceived to move) differ from the representations used for implicit processing of position (i.e. *where* an object is; cf. Brenner & Smeets, 1994; Smeets & Brenner, 1995a) – and involve different cortical areas – or, alternatively, if a shared velocity representation might cause greater differences in perceived speed than in the FLE. The latter explanation could reconcile findings of Vaziri-Pashkam and Cavanagh (2011), as at low luminance the effects of perceived speed did not significantly affect the overall magnitude of the FLE any more than the effects of physical speed.

Furthermore, there was a large variability for individual observers in Experiments 1A and 1B. This could be partly attributed to measurement noise, so we can suspect that the effect of perceived speed (although robust within individuals) might explain only a small proportion of the variance in the illusion across individuals in our study. Other factors might also influence responses in the FLE (thereby contributing to error variance). Kanai et al. (2004) reported FLE biases in the same direction, but they explain their effect of contrast using uncertainty (see also Fu, Shen, & Dan, 2001; Maus & Nijhawan, 2006, 2009; Purushothaman et al., 1998; Vreven & Verghese, 2005). According to this interpretation, increases in FLE magnitude are caused by increases in positional uncertainty, due to decreased contrast rather than increased perceived speed. This reasoning is also compatible with Experiment 1 if the dynamic pattern would increase uncertainty; presumably, dynamically patterned or low contrast objects would be expected to elicit weaker or noisier position signals for the visual system to work with. In turn, the perceived position might depend more strongly on predictions originating from extrapolation processes, and be less strongly influenced by (uncertain) sensory information. This would, in principle, be expected to cause increases in FLE magnitude for dynamic pattern or low contrast conditions as we observe here. However, the hypothesized effects of uncertainty have been shown to have little impact on judgments of speed (Stocker & Simoncelli, 2006), and observers can accurately report the extrapolated positions of moving objects despite decreases in visibility (Graf, Warren, & Maloney, 2005), making this interpretation unlikely.

Instead, we might consider if there is an indirect or additional influence of uncertainty. Uncertainty could predict an increase in error variance, and this would be expected to work against any systemic biases, rather than cause one. This would be compatible with the absence of a cross-effect correlation. Another possible influence on our effects is not in terms of uncertainty, but in the amount of attention allocated to the moving object. We cannot rule this possibility out, as we did not explicitly control or measure attention in our different conditions. Conceivably, an object containing a dynamic pattern might require more attention to localize than an object containing a static texture. If this were true, then we might expect a greater FLE, based on studies showing that the FLE increases when attention is divided (e.g., across concurrent tasks, across multiple stimuli, or for increased speeds; Sarich, Chappell, & Burgess, 2007; Scocchia, Actis-Grosso, de’Sperati, Stucchi, & Baud-Bovy, 2009). This explanation also applies if low contrast requires more attention than high contrast. These alternative interpretations need to be tested by future experiments.

Overall, the pattern of results is consistent with the notion that the magnitude of the FLE depends on a neural representation of velocity that is sensitive to (some of) the same experimental manipulations that also affect over reports of perceived speed. This finding is consistent with an explanation of the FLE in terms of visual motion extrapolation. It also corroborates a growing body of evidence supporting the existence of neural mechanisms involved in extrapolation in both animal models (e.g., Benvenuti et al., 2020; Berry et al.,1999; Jancke, Erlhagen, Schöner, & Dinse, 2004; Palmer, Marre, Berry, & Bialek, 2015; Schwartz, Taylor, Fisher, Harris, & Berry, 2007; Subramaniyan et al., 2018; Sunberg, Fallah, & Reynolds, 2006) and human neuroimaging (e.g., Blom, Feuerriegel, Johnson, Bode, & Hogendoorn, 2020; Ekman, Kok, & de Lange, 2017; Hogendoorn & Burkitt, 2018; Schneider, Marquardt, Sengupta, De Martino, & Goebel, 2019), and is consistent with several decades investigating motion extrapolation in perception (Hogendoorn, 2020; Hubbard, 2005, 2014, 2018; Maus et al., 2010; Nijhawan, 2002, 2008).

## Acknowledgements

This study was supported by an Australian Research Council (ARC) Discovery Projects Grant awarded to H.H. (DP180102268).

Commercial relationships: none.

## Notes

### Competing Interest Statement

The authors have declared no competing interest.

### Summary of Updates

Minor revisions made to clarify

